# PyKappa: Rule-based modeling in Python

**DOI:** 10.64898/2026.09.18.752660

**Authors:** Berk A. Alpay, Amber Lin, August C. Damiani, Walter Fontana

## Abstract

**Summary:** Rule-based languages have proven effective for modeling systems of interacting structured entities as typically encountered in chemistry and molecular biology. We present PyKappa, a rule-based modeling package written in Python whose interpreted nature enables interactive simulation and analysis, including by agentic AI. The package seeks to broaden the base of developers by utilizing a widely known programming language and serves as an easy-to-deploy teaching tool. Using PyKappa, we conduct a case study of phase separation.

**Availability:** The official PyKappa website is https://pykappa.org, where code fo the phase separation analysis can also be found. The PyKappa source code is hosted at https://github.com/berkalpay/pykappa.

---

Many physical and biological systems involve interactions between structured complexes of “atomic” agents. What constitutes an atomic agent depends on the model’s level of description. In chemistry, agents are atoms and complexes are molecules, but for a systems biologist attending to signaling pathways, an atomic agent might be a protein that can noncovalently bind other molecules to form complexes. Interactions between complexes consist in the modification of a *pattern* of agents they contain.

The combinatorial possibilities inherent in pattern-based interaction pose a challenge for modeling system dynamics with techniques, such as Petri nets or differential equations, that require specification of all possible complexes at the outset, as the number of equations required can quickly become unmanageable [1,2]. However, if the structure of complexes can be usefully abstracted from their geometry, they may be represented as graphs amenable to graph-rewriting, a rigorous technique for succinctly specifying rules of graph modification [3]. A *rule* specifies, agent by agent, the transformation of a source graph pattern into a target pattern. Upon identifying a subgraph of a mixture of agents as an instance of the source, a rule instructs the in-place substitution of the subgraph by the target. Because a pattern can occur multiple times in the same complex or in many structurally distinct complexes, rules curtail the combinatorial explosion by only implicitly specifying the possible, like a grammar implicitly specifies all sentences, rather than through explicit enumeration.

Two rule-based languages, BNGL [4,5] and Kappa [2,6], alongside stochastic simulators [7–9] have been developed independently. BNGL has roots in computational biology [10], and Kappa in computer science [11,12]. They are almost identical in syntax and aim to model systems of heterogeneous interactions. Both are founded on the concept of a “site graph.” Unlike a plain graph, the nodes, i.e. agents, in a site graph possess named sites meant to represent interaction resources, such as valences or functional groups of molecules in chemistry, and binding sites or post-translationally modifiable residues of proteins in biology. At any time, a site can have at most one edge, i.e. bond, to a site of another agent. To generate dynamical behavior, a rule is assigned a stochastic rate constant, which is the parameter of an exponential probability density for timing the application of the rule to a chosen match of its source graph in the mixture. The overall propensity with which a rule generates a state transition is determined by its rate constant times the number of embeddings of the source graph in the mixture, which is mass action. A rule-based model can be effectively viewed as a program written in a domain-specific language that is executed stochastically using efficient algorithms [13] adapted to this purpose. Static analysis can be used to predict qualitative aspects of a model’s behavior prior to execution and therefore assist with model-building [14].

In general, rule-based languages are powerful tools for reasoning about mechanisms underlying the dynamics of biomolecular networks, providing a transparent and executable representation of biochemical facts at a level of abstraction often useful in biology. Accompanying its use in molecular biology [2,15], but also synthetic biology [16], disease [17,18], epidemiology [19], and animal behavior [20], these languages have been augmented by an ecology of tools [14,21–27] and theoretical advances [28–35]. Major areas of development have included the visualization of models [24,27], their translation from English [36] and between formal languages [37], including between BNGL and Kappa [23], and formal analysis [14,35]. In Kappa, these tools, including the primary simulator KaSim [6], exist as separate, compiled executables.

To provide an interpreted rule-based modeling environment, we created PyKappa, an open-source Python package for constructing, simulating, and analyzing Kappa models. Extant simulators are written in compiled languages, C++ (BNGL [9]) and OCaml (Kappa [7]), and although they are accordingly fast, an interpreted environment provides several advantages. First is enabling and encouraging community development. Given the still largely untapped potential of rule-based modeling, the tools needed to make, analyze, and execute them would benefit from broader community engagement. A Python code base is an incentive, given that Python has become the lingua franca of scientific computing. Clarity is a core principle of the PyKappa API, making it highly accessible to users and to developers who wish to contribute to the Kappa ecosystem or extend the language and simulator to suit specialized purposes. A second advantage is interactivity. The internal state of compiled simulators can be accessed and modified via prespecified commands, such as with KaSim’s “intervention language”, that specify conditions at which the system state should be transcribed or perturbed. Because it is interpreted, PyKappa obviates the need for such a language altogether; after any update step the state of the simulator and the mixture are available to the modeler for programmatic inspection, analysis and intervention. The interpreted nature of PyKappa thereby also facilitates the use of Jupyter notebooks [38] and the deployment of agentic artificial intelligence in modeling. The third advantage is in teaching. The ease of installation (via the Python Package Index), clear documentation, detailed language reference [39], and the ubiquity of Python make PyKappa an effective resource in the classroom for conveying principles of systems biology through the interactive use of transparent mechanistic models approachable with varying levels of mathematical background. Examples of systems across various fields modeled with PyKappa — such as dynamics of Michaelis-Menten enzyme catalysis, the lac operon gene regulatory system, and vector-borne disease — can be found at the PyKappa website.

PyKappa is object-oriented in design, with a hierarchical structure that mirrors the structure of Kappa models. The core classes for simulation are sites, agents, components, patterns, mixtures, and systems. Sites belong to agents, agents form connected components, components comprise patterns, rules target patterns, and systems apply the transformations specified by rules to the mixture. In addition to the core simulation engine, PyKappa provides tools for analyzing the underlying rule sets as well as the results of stochastic simulation.

PyKappa is compatible with KaSim in that Kappa models can be run with both engines. While the built-in, interpreted PyKappa engine is suitable for development and execution of smaller models, scaling up to large particle numbers may require compiled simulation in KaSim. PyKappa thus provides functionality to forward a mixture to KaSim for simulation and then retrieve intermediate outputs and the new mixture state for continuation within the interpreted PyKappa environment. In the extreme case, in which a model must be simulated entirely with KaSim, PyKappa still considerably expands on the functionality of the extant Python wrapper, kappy, in that it fully represents Kappa objects in Python and provides associated analysis tools.

We illustrate PyKappa with a novel yet simple application of rule-based modeling in molecular systems biology, phase separation. In particular, we model liquid-liquid phase separation, the separation of a homogeneous mixture into two or more distinct parts. Separation usually comes about as a result of multivalent interactions, in biology typically between proteins and nucleic acids or just between proteins [40].

Our model posits two agent types with respective sites, `A(l,r,b)` and `B(d,a1,a2,a3)`. The interactions are defined in terms of five simple reversible rules. For example, the rule A(l[.]), A(r[.]) -> A(l[1]), A(r[1]) asserts that an instance of A can bind at its unoccupied site l another instance of A at its unoccupied site r, regardless of the state at site b. This rule implies polymerization, since it can generate strings or rings of any size, for two strings of A agents so-generated match the left pattern of the rule and can be concatenated. The left pattern could also match two A agents within the same string, leading to intramolecular ring closure. Another rule specifies that B-agents can dimerize on d. Three further rules assert that sites `a1, a2,` and `a3` of `B` can each bind an unoccupied site `b` of an `A`.

To set the stochastic rate constants of each rule, it is perhaps clearest to first define deterministic rate parameters: `A-`agents bind each other weakly with *K*_*d*_ = 10^*−*6^ M, and bonds involving `B-`agents are slightly stronger with *K*_*d*_ = 10^*−*7^ M. Assuming a value for the on-constant, *k*_on_, the respective off-constants are obtained as *K*_*d*_ *k*_on_. The stochastic equivalent of each deterministic rate constant is then *γ*_on_ = *k*_on_*/*(*AV*) where *A* is Avogadro’s constant. Assuming a system volume of *V* = 1 pL (roughly that of a mammalian cell) and concentrations of both agent types of 100 nM translates to 60,220 particles of each agent. There is no need to simulate with so many agents, so we scale the system volume by a factor of *σ* = 0.0075, resulting in only 451 particles of each agent type. The stochastic on-rates are recomputed accordingly.

The rate constants we have issued are independent of whether binding interactions are taking place within the same molecule or between two separate molecules. Molecularity is, however, an important consideration in our model. Binding involves the interaction of two sites with a favorable free-energy contribution from electrostatic interactions and the burying of hydrophobic surfaces. It also involves an unfavorable contribution from the loss of positional entropy [41], Δ*G*_pos_, which is much lower when two sites belong to the same complex within what is effectively a much smaller reaction volume. For the system to be biophysically sensible, in the absence of a spatial representation, we must assign two rate constants to the rule, one for each molecularity of interaction. (In addition to considerations of reaction rates, the distinction between inter- and intramolecular binding behooves simulation engines to efficiently track connected components, as the locality of graph-matching is broken in the presence of this distinction.)

We derive each intramolecular on-rate from the corresponding intermolecular on-rates we have already computed. Specifically, we set the intramolecular on-rate 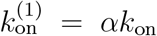 with *α* = exp(Δ*G*_pos_*/RT*). Since positional entropy scales with the logarithm of volume, *α* scales linearly with volume. Thus, 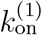 is independent of volume, as for a unimolecular reaction, although it is conceptually still a bimolecular rate constant characterizing the reactive collision between two sites. We set *α* = 6.7 × 10^4^ (Δ*G*_pos_ ≈ 6.5 kcal mol^*−*1^ at 20^*°*^C) and *k*_on_ = 10^8^ s^*−*1^*M* ^*−*1^, which is roughly diffusion-limited. Setting this parameter sets the intramolecular on-rates, and so the rules governing our model are now completely specified.

We perform stochastic simulation to analyze the dynamics of the system, beginning with simulation of a mixture of purely `A-`agents and one of purely B-agents. While `A-`agents can, in principle, form linear and cyclic polymeric strings, the affinity is so weak that an equilibrium mixture of `A` at a concentration of 100 nM consists mostly of monomers and a small fraction of dimers. A pure mixture of B is limited to dimers. However, a mixture of both `A` and `B` permits a `B` to bind up to three `A-`agents slightly more stably than `A-`agents can bind each other, providing a better opportunity for `A-`agents to close mixed cycles. Because cycle formation is intramolecular, it is driven by an on-rate that is higher by a factor of *α*. Cycles permit a structure to withstand more bond losses before breaking apart. Figure 1A indicates that the mixture initially settles into a meta-stable state where larger aggregates form but their cycle density is insufficient to tip any of them towards accretion before bond losses cause shrinkage.

**Figure 1.**
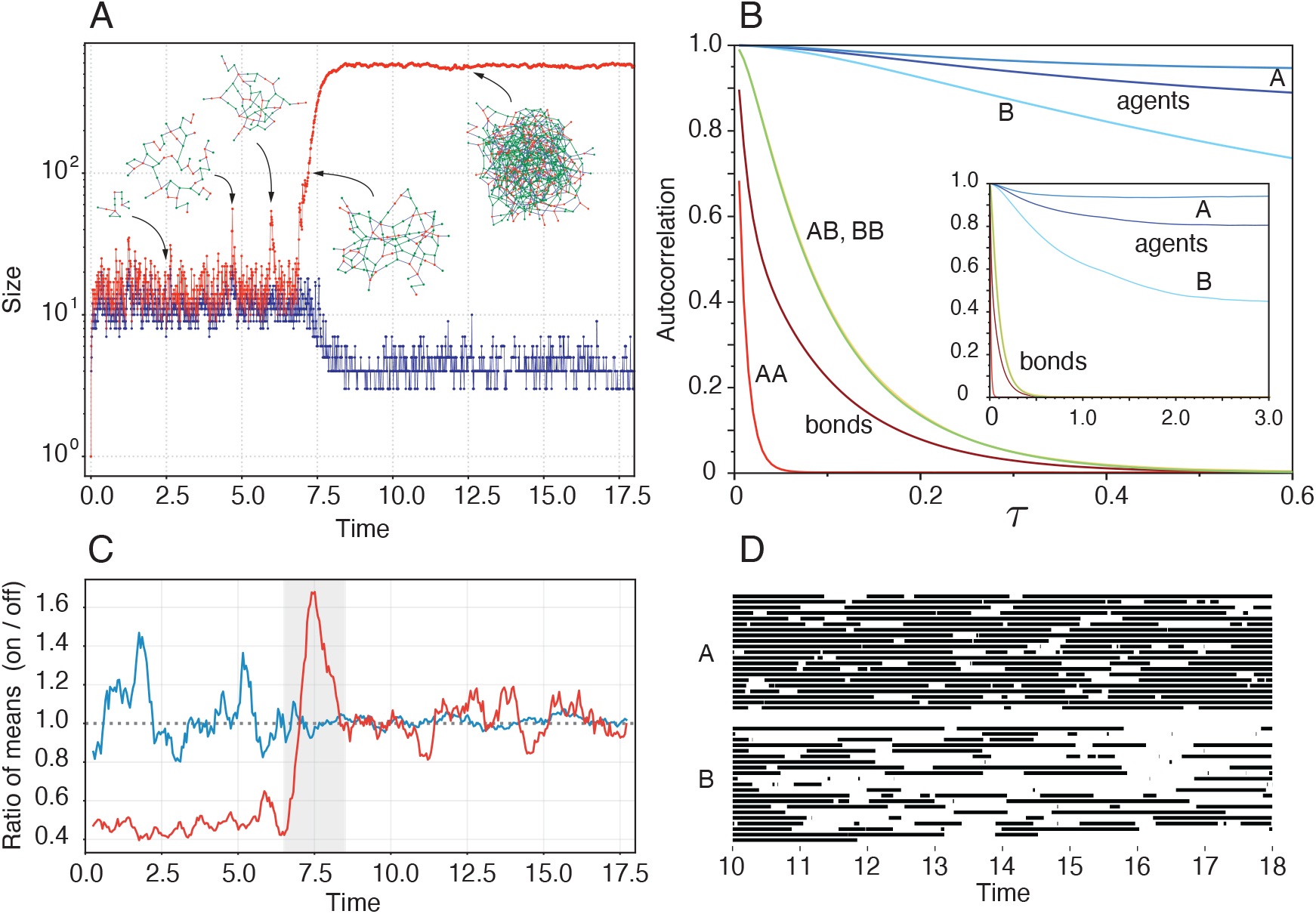
Nucleated phase separation. **A:** Sizes of the largest complex (“maximer”, red) and next-to-largest complex (blue). Prior to condensation small maximers change rapidly; after separation a large complex—a “phase”—emerges whose autocorrelation (**B**) can be written in terms of Jaccard distances *d*(*M*_*t*_, *M*_*t*+*τ*_), *R*(*τ*) = 1 − ⟨*d*(*M*_*t*_, *M*_*t*+*τ*_)^2^⟩, with *M*_*t*_ the phase at time *t* understood as a set of agents or a set of bonds. While the bonds become rapidly uncorrelated, the phase retains long-term memory of agents (inset) because of the small system size. This is illustrated by tracking the presence (black) and absence (white) of 20 agents in the phase (panel **D**). Agents cycle in and out of the phase repeatedly. The phase has a higher avidity (numerous weak binding opportunities) for `A-`agents than `B-`agents. The ratio of `A:B` in the phase at stationarity is 2.1:1. **C:** Analyzing the phase graphs at time *t >* 10 we compute the instantaneous fluxes of bond formation (“on”) and dissociation (“off”), depending on whether they are purely internal, leaving size unchanged, or involve loss or capture of agents (“external”). The mean of each of these four fluxes is computed over a window [*t* − 0.25, *t* + 0.25], *t >* 10, stepping the center every 0.05 time units. Taking the ratio of these means (on/off) yields one ratio for internal (blue) and one for external (red) bond rearrangements. Both end up fluctuating around 1, indicating thermodynamic equilibrium. The internal rearrangements are fluctuating around 1 also in the metastable state, suggesting that the (fictitious) “average structure” over the rapidly changing maximers is in internal bond equilibrium, yet fragile since the gain/loss ratio is tilted towards loss roughly 1:2.

Eventually fluctuations yield a complex with sufficient cycle density to trigger phase separation, which is here the thermodynamically stable state where bond density and growth versus shrinkage attain equilibrium (Figure 1C). The “liquidity” of the new phase is evident from its autocorrelation (Figure 1B): the phase exchanges agents slowly with the remaining mixture, but bonds rearrange rapidly. In this framework, a liquid is a graph whose bond structure is highly dynamic, causing agents to quickly change neighborhood, while maintaining sufficient connectivity to retain agents long enough to establish a bulk phase. Unlike for bonds, the agent autocorrelation does not go to zero because particle numbers are small enough that any given particle is cycled in and out of the liquid phase multiple times, preventing complete loss of memory (Figure 1D). The possible bond rearrangements are much too numerous to recur in any meaningful fraction. Similarity between this system and Axin/APC scaffolding in canonical Wnt signaling is purely coincidental.

## Acknowledgments

This work was funded by the John Templeton Foundation under grant ID 63448 and a Harvard PRISE fellowship to A.C.D. The opinions expressed in this work are those of the authors and do not necessarily reflect the views of the John Templeton Foundation.

## References

1. Hlavacek, W. S., Faeder, J. R., Blinov, M. L., Perelson, A. S. & Goldstein, B. The complexity of complexes in signal transduction. Biotechnology and Bioengineering 84, 783–794 (2003).

2. Danos, V., Feret, J., Fontana, W., Harmer, R. & Krivine, J. Rule-based modelling of cellular signalling in International Conference on Concurrency Theory (2007), 17–41.

3. Ehrig, H., Kreowski, H.-J., Maggiolo-Schettini, A., Rosen, B. K. & Winkowski, J. Transformations of structures: an algebraic approach. Mathematical Systems Theory 14, 305–334 (1981).

4. Faeder, J. R., Blinov, M. L., Goldstein, B. & Hlavacek, W. S. Rule-based modeling of biochemical networks. Complexity 10, 22–41 (2005).

5. Harris, L. A. et al. BioNetGen 2.2: advances in rule-based modeling. Bioinformatics 32, 3366–3368 (2016).

6. Boutillier, P. et al. The Kappa platform for rule-based modeling. Bioinformatics 34, i583–i592 (2018).

7. Danos, V., Feret, J., Fontana, W. & Krivine, J. Scalable simulation of cellular signaling networks in Asian Symposium on Programming Languages and Systems (2007), 139–157.

8. Yang, J., Monine, M. I., Faeder, J. R. & Hlavacek, W. S. Kinetic Monte Carlo method for rule-based modeling of biochemical networks. Physical Review E 78, 031910 (2008).

9. Sneddon, M. W., Faeder, J. R. & Emonet, T. Efficient modeling, simulation and coarse-graining of biological complexity with NFsim. Nature Methods 8, 177–183 (2011).

10. Hlavacek, W. S. & Faeder, J. R. The complexity of cell signaling and the need for a new mechanics. Science Signaling 2, pe46–pe46 (2009).

11. Danos, V. & Laneve, C. Formal molecular biology. Theoretical Computer Science 325, 69–110 (2004).

12. Fontana, W. Systems biology, models, and concurrency. ACM SIGPLAN Notices 43, 1–2 (2008).

13. Gillespie, D. T. Exact stochastic simulation of coupled chemical reactions. The Journal of Physical Chemistry 81, 2340–2361 (1977).

14. Boutillier, P. et al. KaSa: A static analyzer for Kappa in International Conference on Computational Methods in Systems Biology (2018), 285–291.

15. Chylek, L. A., Harris, L. A., Faeder, J. R. & Hlavacek, W. S. Modeling for (physical) biologists: an introduction to the rule-based approach. Physical Biology 12, 045007 (2015).

16. Wilson-Kanamori, J., Danos, V., Thomson, T. & Honorato-Zimmer, R. Kappa rule-based modeling in synthetic biology in Computational Methods in Synthetic Biology 105–135 (Springer, 2014).

17. Bougueon, M. et al. A rule-based multiscale model of hepatic stellate cell plasticity: Critical role of the inactivation loop in fibrosis progression. PLoS Computational Biology 20, e1011858 (2024).

18. Larkin, C. I., Dunn, M. D., Shoemaker, J. E., Klimstra, W. B. & Faeder, J. R. A detailed kinetic model of Eastern equine encephalitis virus replication in a susceptible host cell. PLoS Computational Biology 21, e1013082 (2025).

19. Waites, W., Cavaliere, M., Manheim, D., Panovska-Griffiths, J. & Danos, V. Rule-based epidemic models. Journal of Theoretical Biology 530, 110851 (2021).

20. Bouguéon, M., Petrov, T. & Salazar, A. A rule-based modeling approach for studying animal collectives: A case study of juvenile honeybee thermotaxis in International Conference on Computational Methods in Systems Biology (2025), 174–194.

21. Hu, B., Matthew Fricke, G., Faeder, J. R., Posner, R. G. & Hlavacek, W. S. GetBonNie for building, analyzing and sharing rule-based models. Bioinformatics 25, 1457–1460 (2009).

22. Lopez, C. F., Muhlich, J. L., Bachman, J. A. & Sorger, P. K. Programming biological models in Python using PySB. Molecular Systems Biology 9, 646 (2013).

23. Suderman, R. & Hlavacek, W. S. TRuML: a translator for rule-based modeling languages in Proceedings of the 8th ACM International Conference on Bioinformatics, Computational Biology, and Health Informatics (2017), 372–377.

24. Sekar, J. A. P.Tapia, J.-J. & Faeder, J. R. Automated visualization of rule-based models. PLoS Computational Biology 13, e1005857 (2017).

25. Suderman, R., Fricke, G. M. & Hlavacek, W. S. Using RuleBuilder to graphically define and visualize BioNetGen-language patterns and reaction rules in Modeling Biomolecular Site Dynamics: Methods and Protocols 33–42 (Springer, 2019).

26. Camporesi, F., Feret, J. & Lý, K. Q. KADE: A tool to compile Kappa rules into (reduced) ODE models in International Conference on Computational Methods in Systems Biology (2017), 291–299.

27. Liguori-Bills, N. & Blinov, M. L. bnglViz: online visualization of rule-based models. Bioinformatics 40, btae351 (2024).

28. Danos, V., Feret, J., Fontana, W., Harmer, R. & Krivine, J. Rule-based modelling, symmetries, refinements in International Workshop on Formal Methods in Systems Biology (2008), 103–122.

29. Danos, V., Feret, J., Fontana, W. & Krivine, J. Abstract interpretation of cellular signalling networks in International Workshop on Verification, Model Checking, and Abstract Interpretation (2008), 83–97.

30. Feret, J., Danos, V., Krivine, J., Harmer, R. & Fontana, W. Internal coarse-graining of molecular systems. Proceedings of the National Academy of Sciences 106, 6453–6458 (2009).

31. Hogg, J. S., Harris, L. A., Stover, L. J., Nair, N. S. & Faeder, J. R. Exact hybrid particle/population simulation of rule-based models of biochemical systems. PLoS Computational Biology 10, e1003544 (2014).

32. Harmer, R., Danos, V., Feret, J., Krivine, J. & Fontana, W. Intrinsic information carriers in combinatorial dynamical systems. Chaos 20, 037108 (2010).

33. Danos, V. et al. Graphs, rewriting and pathway reconstruction for rule-based models in FSTTCS 2012 18 (2012), 276–288.

34. Danos, V., Harmer, R. & Honorato-Zimmer, R. Thermodynamic graph-rewriting. Logical Methods in Computer Science 11 (2015).

35. Laurent, J., Yang, J. & Fontana, W. Counterfactual resimulation for causal analysis of rule-based models in IJCAI (2018), 1882–1890.

36. Gyori, B. M. et al. From word models to executable models of signaling networks using automated assembly. Molecular systems biology 13, MSB177651 (2017).

37. Tapia, J.-J. & Faeder, J. R. The Atomizer: extracting implicit molecular structure from reaction network models in Proceedings of the International Conference on Bioinformatics, Computational Biology and Biomedical Informatics (2013), 726–727.

38. Kluyver, T. et al. Jupyter Notebooks–a publishing format for reproducible computational workflows in Positioning and Power in Academic Publishing: Players, Agents and Agendas 87–90 (IOS press, 2016).

39. Fontana, W., Boutillier, P., Feret, J. & Krivine, J. The Kappa Language and Kappa Tools: A User Manual and Guide Version 4 (Aug. 2026). https://kappalanguage.org/static/manual.pdf.

40. Banani, S. F., Lee, H. O., Hyman, A. A. & Rosen, M. K. Biomolecular condensates: organizers of cellular biochemistry. Nature Reviews Molecular Cell Biology 18, 285–298 (2017).

41. Saiz, L. & Vilar, J. M. Stochastic dynamics of macromolecular-assembly networks. Molecular Systems Biology 2 (2006).

